# Structural insights into the high selectivity of the anti-diabetic drug mitiglinide

**DOI:** 10.1101/2022.04.26.489624

**Authors:** Mengmeng Wang, Jing-Xiang Wu, Lei Chen

## Abstract

Mitiglinide is a highly selective fast-acting anti-diabetic drug that inhibits pancreatic K_ATP_ channels to induce insulin secretion. However, how mitiglinide binds K_ATP_ channels remains unknown. Here, we show the cryo-EM structure of the SUR1 subunit in complex with mitiglinide. The structure reveals that mitiglinide binds inside the common insulin secretagogue-binding site in the transmembrane domain of SUR1, locking SUR1 in a NBD-separated inward-facing conformation. The detailed structural analysis uncovers the molecular basis of the high selectivity of mitiglinide.

## Introduction

More than 400 million people are living with diabetes worldwide and type 2 diabetes (T2DM) accounts for nearly 90% of diabetes patients (Chatterjee et al., 2017). Dysfunction of insulin secretion is one of the hallmarks of T2DM (Daryabor et al., 2020). Previous studies have established that K_ATP_ channel plays an essential role in insulin secretion and is a drug target for diabetes (Gribble and Reimann, 2003).

K_ATP_ channel is a hetero-octamer complex composed of four pore-forming Kir6 (kir6.1 or Kir6.2) subunits and four regulatory sulfonylureas receptor subunits (SUR1 or SUR2). K_ATP_ channels can sense intracellular ATP/ADP ratio using the inhibitory ATP binding site on Kir6 and the activating Mg-ADP binding site on SUR, coupling the metabolism status of the cell to its membrane potential (Ashcroft and Gribble, 1998). K_ATP_ channels are widely distributed in many tissues, including brain (Ashford et al., 1988), muscles (Noma, 1983; Spruce et al., 1985), and endocrine cells (Cook and Hales, 1984), and play important physiological functions. In pancreatic β-cells, K_ATP_ channels are mainly formed by Kir6.2 and SUR1 and play key roles in controlling insulin secretion (Ashcroft and Rorsman, 1989). When blood glucose level increases, the intracellular ATP/ADP ratio also increases accordingly, leading to the suppression of the K_ATP_ channel activity. The inhibited potassium efflux through the K_ATP_ channel induces depolarization of the β-cell membrane, resulting in the subsequent activation of voltage-gated calcium channel, calcium influx, and insulin secretion (Nichols, 2006; Rorsman and Trube, 1985). Mutations in genes encoding either Kir6.2 or SUR1 cause disorders in insulin secretion, such as congenital hyperinsulinemia and neonatal diabetes (Ashcroft, 2005). Small molecule drugs that inhibit the pancreatic K_ATP_ channel are widely used to boost insulin secretion for the treatment of diabetes and are therefore named insulin secretagogues (IS) (Gribble and Reimann, 2003).

ISs are chemically diverse small molecules, including sulphonylureas, such as glibenclamide (GBM), and glinides, such as repaglinide (RPG) and mitiglinide (MTG) (Wu et al., 2020). ISs bind to the SUR subunits to inhibit the K_ATP_ channels(Wu et al., 2020). MTG ((+)-monocalcium bis [(2S)-2-benzyl-3-(cis-hexahydro-2-isoindolinyl carbonyl)propionate]dihydrate), also named KAD-1229, is a glinide that was developed for the treatment of postprandial hyperglycemia (Pratley et al., 2001) and has been approved for the treatment of patients with T2DM in Japan (brand name, Glufast, approved in 2004). MTG has an immediate and short-lasting effect on hypoglycemic action in the postprandial glucose-load state in clinical trials (Phillippe and Wargo, 2013). In vitro experiments supported that MTG increases insulin release from a pancreatic β-cell line MIN6 cell (Mogami et al., 1994; Reimann et al., 2001). The insulin responses to chronic MTG treatment were comparable to chronic RPG or GBM treatment in MIN6 cells (Reimann et al., 2001). MTG can displace [^3^H]-GBM binding to HIT-15 cells with an IC_50_ of 13 nM (Ohnota et al., 1994), suggesting that they might share an overlapped binding site. Notably, MTG is highly selective towards SUR1 over SUR2 (Reimann et al., 2001; Sunaga et al., 2001), and GBM is moderately selective, while RPG is non-selective (Quast et al., 2004). Recent structural studies have uncovered the binding modes of GBM and RPG(Wu et al., 2020), but the exact binding mode of MTG on SUR1 remains unknown. Here, we present the cryo-EM structure of the SUR1 subunit complexed with MTG, allowing the direct visualization of how MTG binds SUR1.

## Methods

### Cell lines

FreeStyle 293-F (Thermo Fisher Scientific) suspension cells were cultured in SMM 293-TI (Sino Biological Inc.) supplemented with 1% fetal bovine serum (FBS) at 37°C, with 6% CO_2_ and 70% humidity. Sf9 insect (Thermo Fisher Scientific) suspension cells were cultured in Sf-900 III SFM medium (Thermo Fisher Scientific) at 27°C.

### Construct of NGFP_linker_SUR1core

We made a truncated *Mesocricetus auratus* SUR1 construct maSUR1core (from 208 to C terminal) based on the previous work (Ding et al., 2019a). We added an N-terminal GFP and MBP tag, a PreScission protease cleavage site, KNtp of mmKir6.2 (Ding et al., 2019a), and a GS rich linker before SUR1core. The construct was made in a modified BacMam vector (Li et al., 2017).

### Electrophysiology

Wild-type SUR1 or its mutants together with CGFP-tagged Kir6.2 were transfected into FreeStyle 293F suspension cells using polyethyleneimine at the cell density of 1.0×10^6^ cells/ml. Cells were cultured in FreeStyle expression medium supplemented with 1% FBS for 24 h and then seeded into the 12 mm dishes for adhesion before recording. Macroscopic currents were recorded in the inside-out mode at +60 mV by Axon-patch 200B amplifier (Axon Instruments, USA). Patch electrodes were pulled by a horizontal microelectrode puller (P-1000, Sutter Instrument Co, USA) to 2.0-3.0 MΩ resistance. Both pipette and bath solution was based on KINT buffer, containing (mM): 140 KCl, 1 EGTA, and 10 HEPES (pH 7.4, KOH). For mitiglinide (Targetmol) and RPG (Abcam) inhibition, both the stock solutions (10 mM mitiglinide and 100 mM RPG) were dissolved in DMSO, and stored at −20°C, and diluted into KINT buffer to the final concentrations before recording. ATP (Sigma) and ADP (Sigma) stocks were prepared on ice, and stored at −20°C. ATP and ADP were dissolved in water and adjusted to pH = 7 by KOH (Sigma). The nucleotide concentration was determined by its extinction coefficient and absorption at 259 nm. Signals were acquired at 5 kHz and low-pass filtered at 1 kHz. Data were further analyzed by pclampfit 10.0 software.

### Protein expression and purification

SUR1core was expressed as described previously (Ding et al., 2019a) and the purification process was carried out with minor modifications. For protein purification, membrane pellets were homogenized in TBS (20 mM Tris-HCl, pH 7.5, 150 mM NaCl) and solubilized in TBS with 1% GDN (Anatrace), supplemented with protease inhibitors (1 mg/ml Leupeptin, 1 mg/ml Pepstatin, 1 mg/ml Aprotinin and 1 mM PMSF), 1mM EDTA and 1mM EGTA for 30 min at 4°C. Unsolubilized materials were discarded after centrifugation at 100,000 g for 30 min and the supernatant was loaded onto a 5 mL Streptactin 4FF (Smart Lifesciences) packed column. Strep column was washed by TBS buffer supplemented with 0.01% GDN, protease inhibitors (1 mg/ml Leupeptin, 1 mg/ml Pepstatin, 1 mg/ml Aprotinin), 1mM EDTA, and 1mM EGTA. Then the column was washed by TBS supplemented with 0.01% GDN, 3mM MgCl_2_, and 1mM ATP. The last washing step buffer was TBS supplemented with 0.01% GDN. The SUR1_core_ was eluted with TBS (50 mM Tris-HCl, pH 7.5, 150 mM NaCl) supplemented with 0.006% GDN and 5 mM desthiobiotin. The eluate was concentrated and supplemented with PreScission protease overnight. The next day, SUR1core was further purified using Superose 6 increase (GE Healthcare) column running with TBS supplemented with 0.006% GDN. Peak fractions were collected and concentrated to A_280_=10 (50 μM). The purified protein was used for cryo-EM sample preparation.

### Cryo-EM sample preparation

100 µM mitiglinide was added to the sample and incubate for 20 min before centrifugation at 40,000 rpm for 30 min using a TLA55 rotor (Beckman). Cryo-EM grids were prepared with Vitrobot Mark IV (FEI) and GIG R0.6/1 holey carbon grids, which were glow-discharged for 120 s using air before adding the protein sample. A 2.5 µl sample was applied to the glow-discharged grid and then the grid was blotted at blotting force at level 2 for 2 s at 100% humidity and 10°C, before plunge-frozen into the liquid ethane.

### Cryo-EM data acquisition

Cryo-grids were screened on a Talos Arctica microscope (Thermo Fisher Scientific) operated at 200 kV. Grids of good quality were further loaded onto Titan Krios microscope (Thermo Fisher Scientific) operated at 300 kV for data collection. Images were collected using a K2 camera (Gatan) mounted post a Quantum energy filter with 20 eV slit and operated under super-resolution mode with a pixel size of 1.045 Å at the object plane, controlled by software Serial EM. Defocus values were set from −1.5 μm to −1.8 μm for data collection. The dose rate on the detector was 8.078 e^-^ pixel^-1^ s^-1^ and the total exposure was 54.3 e^-^A^-2^. Each 6.72 s movie was dose-fractioned into 32 frames.

### Image processing

1,181 Movies were collected and the movies were exposure-filtered, gain-corrected, motion-corrected, mag-distortion-corrected, and binned with MotionCor2 (Zheng et al., 2017), producing dose-weighted and summed micrographs. CTF models of dose-weighted micrographs were determined using Gctf (Zhang, 2016). Gautomatch (developed by Kai Zhang, MRC-LMB) was used for auto-picking. Data processing was initially performed using Relion_3.0 (Zivanov et al., 2018). Particles were extracted from dose-weighted micrographs. After 2D classification (379K particles) and 3D classification with C1 symmetry, 130K particles with good transmembrane domain (TMD) densities were re-extracted and re-centered. In this stage, the mitiglinide density can be observed. The remaining particles were subjected to focused 3D classification with a TMD mask to select around 20K particles (seed particles) with good mitiglinide density. Then the 379K particles were subjected to seed-facilitated 3D classification (Wang et al., 2021) to produce 130K particles in CryoSPARC2 (Punjani et al., 2017). These particles were subjected to non-uniform refinement, CTF refinement, and local non-uniform refinement in cryoSPARC2 to generate a high-resolution map.

### Model building

SUR1 ABC transporter domain in the previously reported K_ATP_ model (PDB ID: 6BJ1) was docked into the density map using UCSF Chimera (Pettersen et al., 2004). The model was manually rebuilt in Coot (Emsley et al., 2010) and refined against the density map using Phenix (Afonine et al., 2018). Figures were prepared using Pymol and UCSF Chimera X (Pettersen et al., 2020).

### Quantification and statistical analysis

Global resolution estimations of cryo-EM density maps are based on the 0.143 Fourier Shell Correlation criterion (Rosenthal and Henderson, 2003). Electrophysiological data reported were analyzed with pclampfit 10.0 software. The number of biological replicates (N) and the relevant statistical parameters for each experiment (such as mean or standard error) are described in figure legends. No statistical methods were used to pre-determine sample sizes.

## Results

### Structure determination of SUR1core in complex with mitiglinide

Previous cryo-EM studies on K_ATP_ channels in complex with GBM (Martin et al., 2017; Wu et al., 2018) or RPG (Ding et al., 2019b) have established that the ABC transporter region of the SUR subunit harbors the IS binding site. Therefore, we used a truncated construct (SUR1core) that encompasses 208-1582 of SUR1, including TMD1, NBD1, TMD2, and NBD2 (Figs. 1a and S1), for subsequent structure determination. Purified SUR1core protein was supplemented with ATP and MTG for cryo-EM sample preparation and data collection. Image processing yielded a reconstruction with an overall resolution of 3.27 Å (Fig. 1b-c, S1-2 and Table S1). High local map quality allows the unambiguous identification of the MTG molecule (Fig 1d-g).

**Fig 1.**
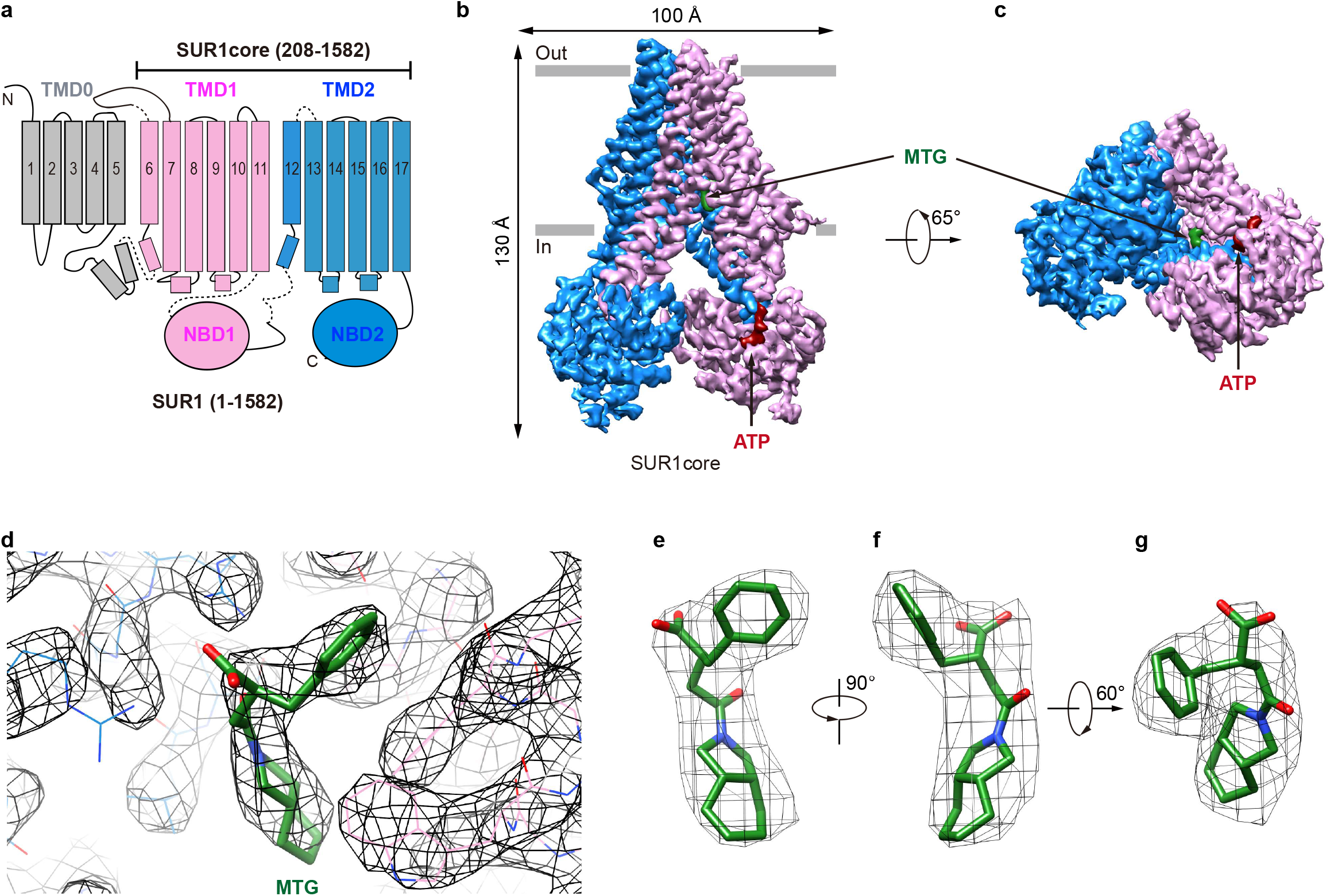
Structure of SUR1 in complex with MTG. **a**, Topology of SUR1 subunits. TMD, transmembrane domain; NBD, nucleotide-binding domain. Transmembrane helices are shown as cylinders. TMD0 fragment, TMD1-NBD1, and TMD2-NBD2 are shown in grey, pink, and blue, respectively. **b**, Side view of the cryo-EM density of the SUR1core. ATP and MTG are shown in red and green, respectively. **c**, Bottom view of the cryo-EM density of the SUR1core. **d**, Densities of MTG and its surrounding amino acids. **e-g**, MTG densities view from different angles.

### The binding mode of mitiglinide

In the presence of ATP and MTG, SUR1core shows an inward-facing conformation with its central vestibule widely open to the cytosol (Fig. 1b-c). ATP binds to the NBD1 and MTG binds inside a pocket in the central vestibule of SUR1 (Fig. 1b-d). The MTG-binding pocket is formed by residues on TM7 and TM8 of TMD1 and TM16 and TM17 of TMD2 (Fig. 2a-b). The interactions between MTG and SUR1 involve both polar interactions and hydrophobic interactions (Fig. 2a-b). The central negatively charged carboxyl group of MTG makes electrostatic interactions with positively charged R1246 on TM16 and R1300 on TM17 (Fig. 2a-b). The benzene ring of MTG stacks with the phenyl group of F433 (Fig. 2a-b). The bulky hexahydro-2-isoindoline group binds in a hydrophobic pocket surrounded by L1241 and T1242 of TMD2 and I381, I385, F433, and W430 of TMD1 (Fig. 2a-b). To understand the role of R1246 and R1300 on the inhibitory function of MTG, we mutated them into alanines individually. We found that both R1246A and R1300A mutations significantly decreased the inhibitory effect of MTG (Figs. 2d and S3), emphasizing their importance on MTG binding and inhibition.

**Fig 2.**
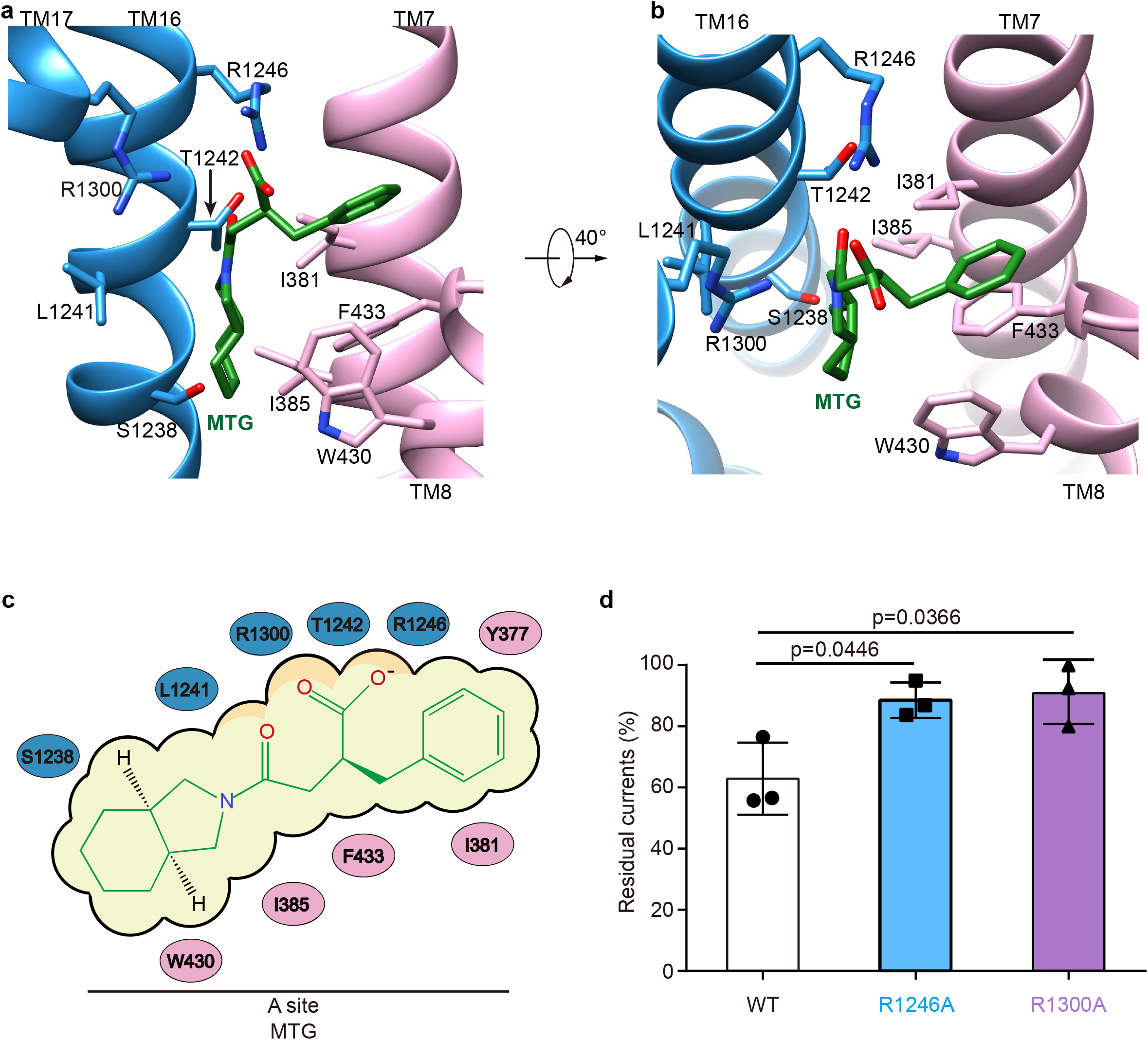
The MTG binding pocket in SUR1. **a-b**, Close-up views of the MTG-binding site. TMD1, TMD2 and MTG are shown in pink, blue, and green, respectively. **c**, Cartoon representation of the interaction between MTG and SUR1. **d**, Wild-type, and mutated K_ATP_ channels inhibition by 10 μM MTG. Data is shown as mean ± SD and n=3 independent patches. p-value was calculated by unpaired two-tailed t-test and indicated above.

### Mitiglinide binds the A-site of SUR1

Accumulated studies on the structure-activity relationship of IS indicate that there are two overlapping binding sites for IS on SUR, the A-site and the B-site (Yan et al., 2006). GBM binds in both the A- and the B-site (Wu et al., 2018), while RPG binds in the B-site (Ding et al., 2019a; Martin et al., 2019). Our structure shows that MTG mainly binds in the A-site (Fig. 3a-c). MTG-interacting residues are almost identical between SUR1 and SUR2, except that T1242 is replaced by a serine in SUR2 and S1238 is replaced by tyrosine in SUR2 (Fig. 3d). Notably, S1238 is near the A-site (Fig. 3a-b), and previous studies found that S1238 is the key determinant for IS selectivity (Dabrowski et al., 2001). In agreement with this, the hexahydro-2-isoindoline group of MTG is near the S1238, and the replacement of S1238 with a tyrosine would likely generate a sterical clash with MTG, abolishing its binding (Fig. 3c).

**Fig 3.**
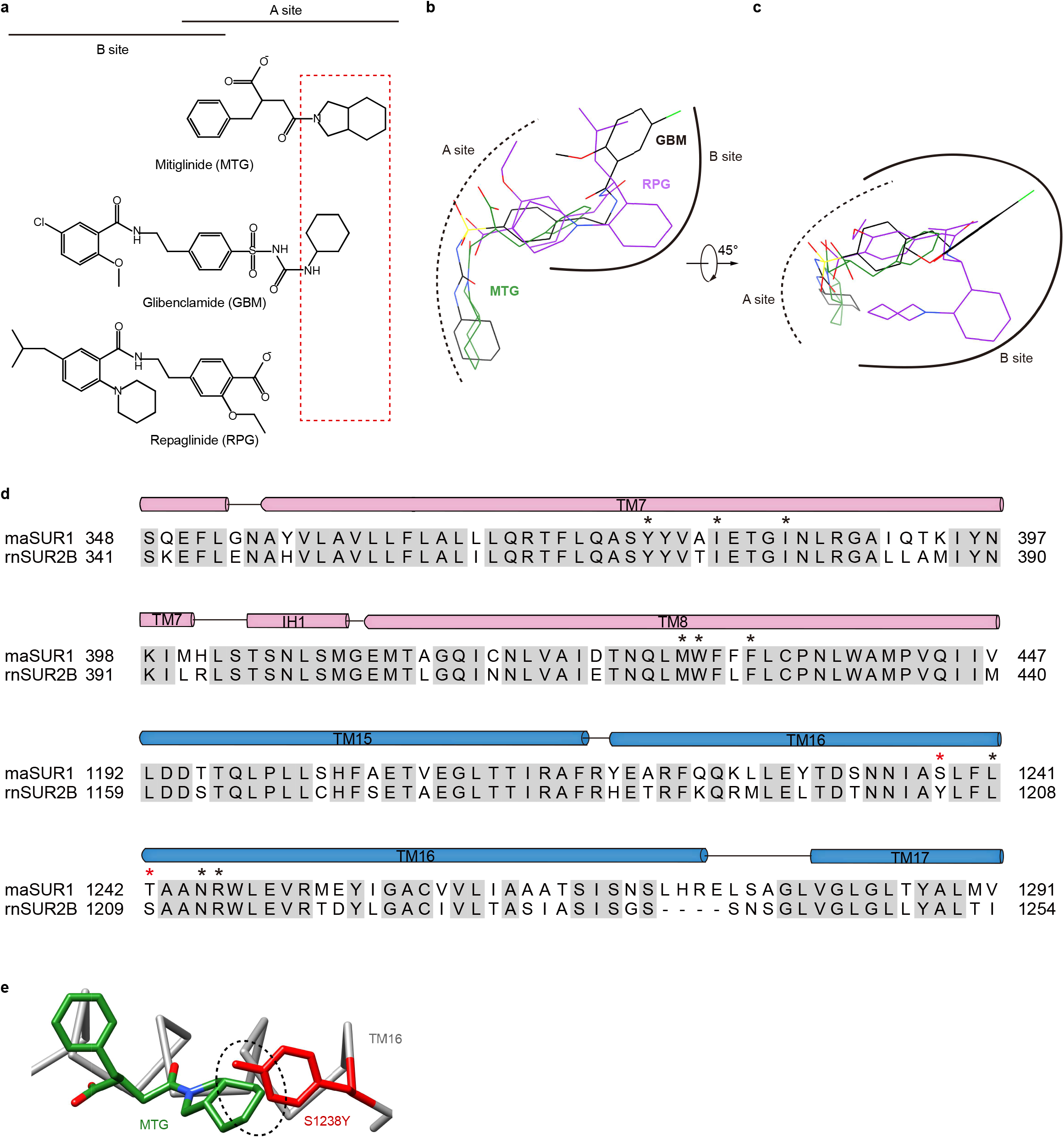
MTG binds in the A-site of SUR1. **a**, Chemical structures of MTG (A-site ligand), RPG (B-site ligand), and GBM (A+B site ligand). The “SUR Isotype Selectivity Determinative Moiety” (SISDM) is boxed in red. **b-c**, Structural superposition of MTG, RPG (PDB ID: 6BJ3) and GBM (PDB ID: 6BAA) in their binding poses. **d**, Sequence alignment of MTG binding pocket between *Mesocricetus auratus* SUR1 (maSUR1) and *Rattus norvegicus* (rnSUR2B). The identical and different MTG-interacting residues are highlighted by black and red asterisks above. **e**, The sterical clashes between MTG (green) and S1238Y (red). The dashed circle highlights clashes.

## Discussion

The structures of SUR1 in complex with IS are now available for RPG, GBM, and MTG. Among them, MTG has the highest selectivity towards SUR1 over SUR2 (Quast et al., 2004). Notably, SUR isotype-selectivity is highly clinically relevant, because SUR1-containing K_ATP_ channels are mainly distributed in pancreatic endocrine cells while SUR2-containing K_ATP_ channels have broad distribution in cardiac muscles and vascular smooth muscles, participating in several key physiological processes in the cardiovascular system, such as the preservation of cardio-protection under ischemic conditions. Moreover, patients with diabetes often have cardiovascular diseases, such as coronary heart disease. Therefore, it is important to consider the off-target side effect of IS treatment for diabetes. The high selectivity of MTG (1,000×) might mitigate the underlying side effects of inhibiting SUR2-containing K_ATP_ channels. In agreement with this theory, it was reported that MTG could preserve the cardio-protective effect of ischemic preconditioning while GBM could not (Ogawa et al., 2007), emphasizing the potential benefit of using highly selective IS for diabetes. In addition, the high selectivity of MTG is also explored to design ^18^F-labeled positron emission tomography tracers for the measurement of β-cell mass in vivo during the progression of diabetes, aiming to better understand the pathogenesis, to facilitate early diagnosis, and to develop novel therapeutics for diabetes (Kimura et al., 2014). Despite the early observation of high selectivity of MTG, how this is achieved was not well-understood previously. Our current structure of SUR1 in complex with MTG shows that the hexahydro-2-isoindoline group is close to S1238, the key residue that is different between SUR1 and SUR2. Therefore, we designate this large hexahydro-2-isoindoline as the “SUR Isotype Selectivity Determinative Moiety” (SISDM). Furthermore, structural comparisons among RPG, GBM, and MTG reveal that the size of SISDM correlates well with their selectivity: larger SISDM confers better SUR-isotype selectivity. Therefore, our structure provides mechanistic insight into the high selectivity of MTG and paves the way for further development of next-generation IS with high selectivity.

## Author Contributions

Lei Chen initiated the project. Mengmeng Wang purified protein, prepared the cryo-EM sample, and performed electrophysiology experiments. Mengmeng Wang and Jing-Xiang Wu collected the data. Mengmeng Wang processed the data, and built and refined the model with the help of Lei Chen. All authors contributed to the manuscript preparation.

## Acknowledgments

Cryo-EM data collection was supported by the Electron microscopy laboratory and Cryo-EM platform of Peking University with the assistance of Xuemei Li, Zhenxi Guo, Bo Shao, Xia Pei, and Guopeng Wang. Part of the structural computation was also performed on the Computing Platform of the Center for Life Science and High-performance Computing Platform of Peking University. We thank the National Center for Protein Sciences at Peking University in Beijing, China for assistance with negative stain EM.

## Data Availability Statement

Atomic coordinates and cryo-EM maps are deposited in EMDB and PDB as EMD-32535 (https://www.ebi.ac.uk/emdb/EMD-32535) and PDB 7WIT (https://www.wwpdb.org/pdb?id=pdb_00007wit);

## Supplementary Material

**Supplementary Figure 1.**
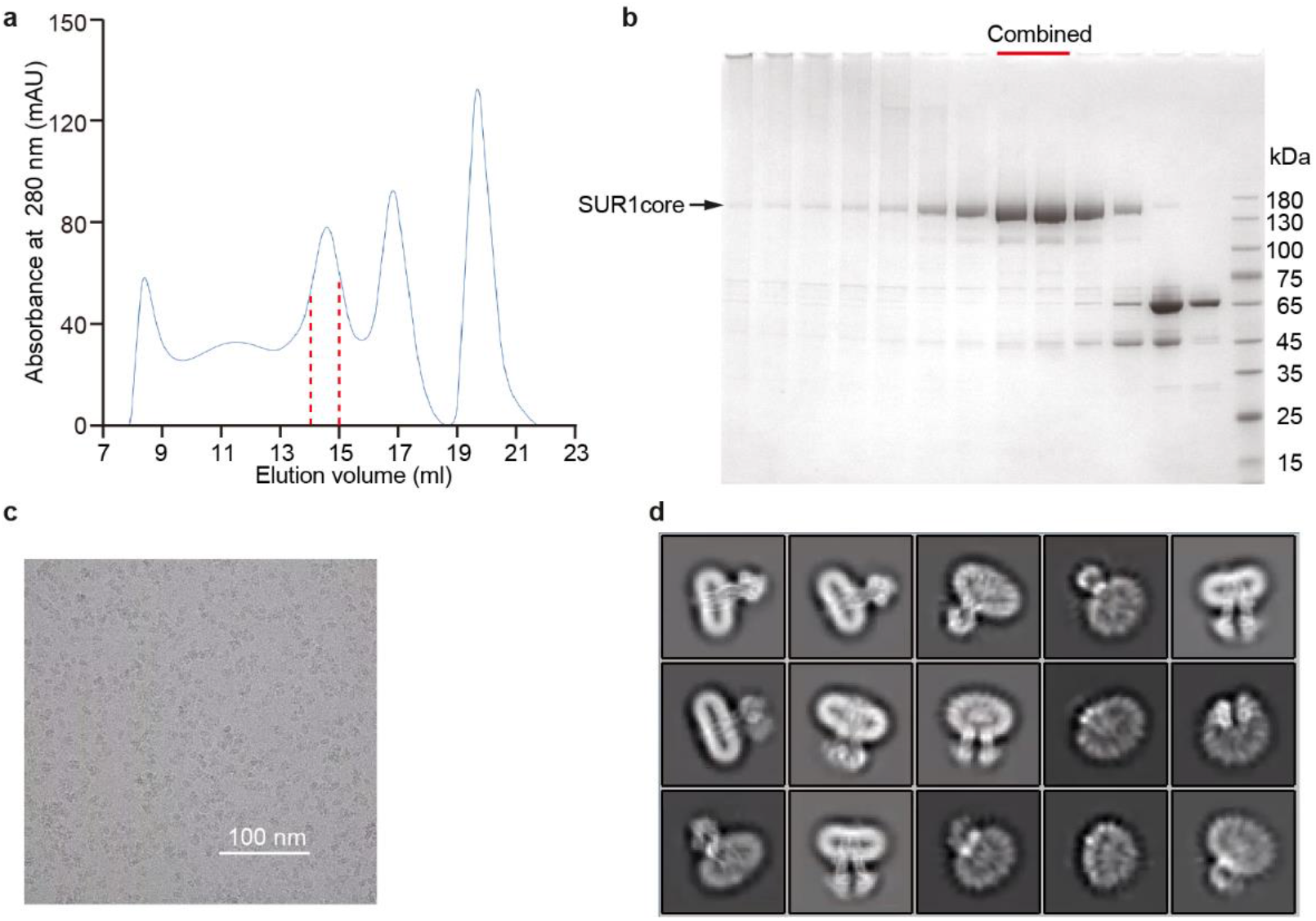
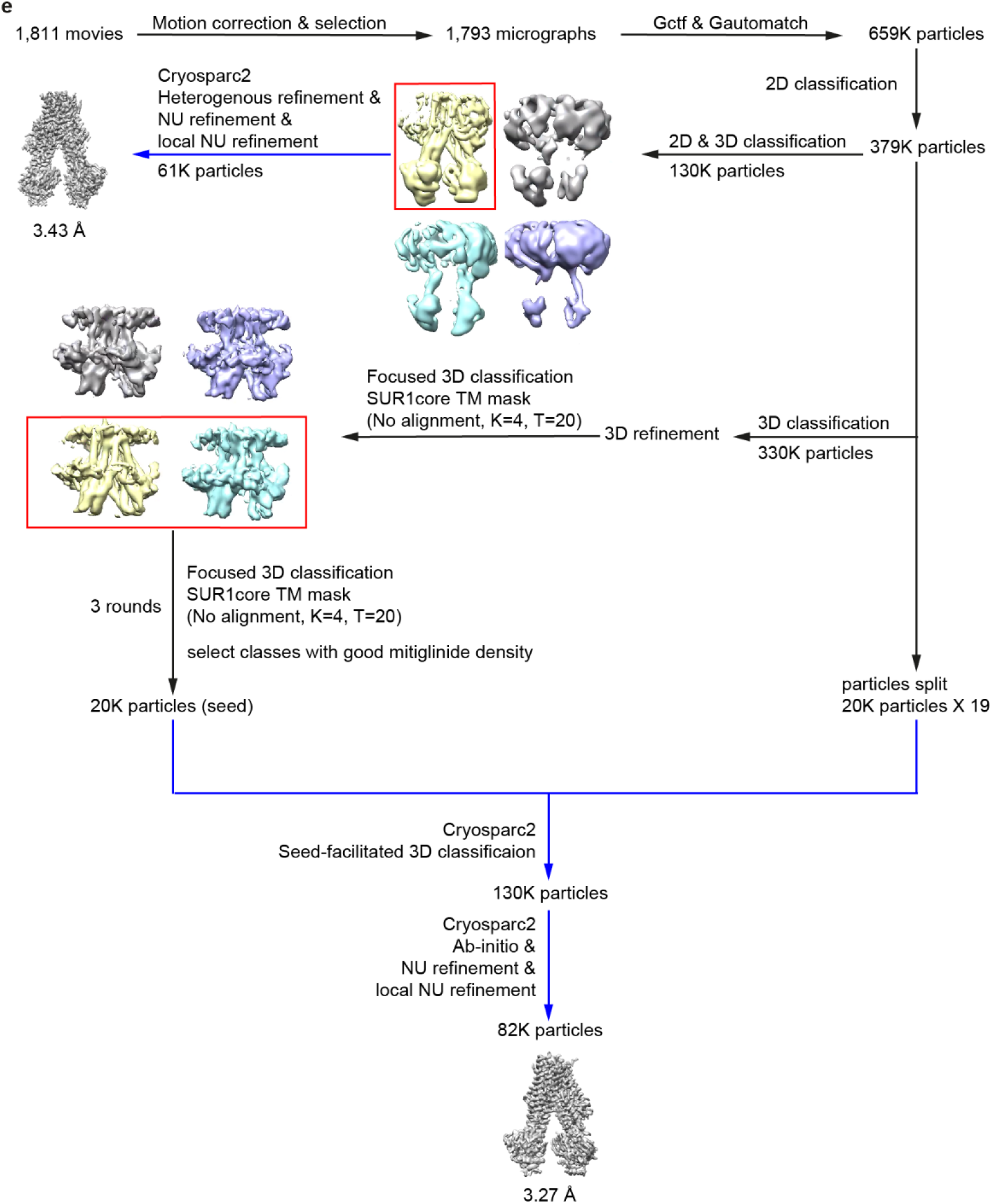
Cryo-EM sample preparation and data processing flow chart. **a**, Size-exclusion chromatography (SEC) elution profile of SUR1core. Fractions between the dashed lines were used for cryo-EM sample preparation. **b**, SDS-PAGE detection of SUR1core. Fractions labeled by red bars were collected for cryo-EM sample preparation. **c**, Representative raw micrograph. **d**, Representative 2D class averages. **e**, Flow chart of cryo-EM image processing of SUR1core in complex with RPG.

**Supplementary Figure 2.**
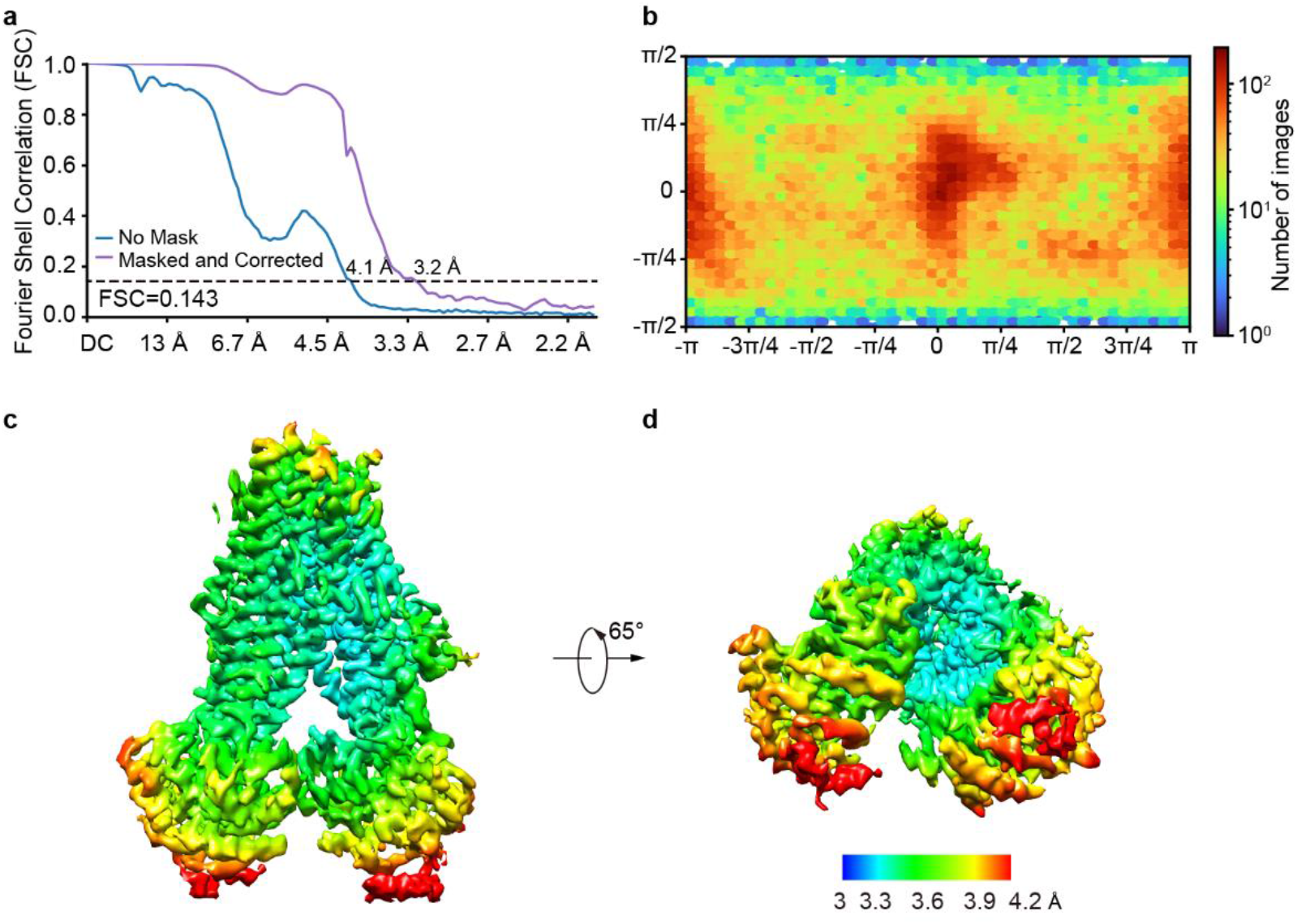
Parameters for the final refined SUR1core Cryo-EM map. **a**, FSC curve of the final refined map is shown. **b**, Angular distribution of SUR1core particles for the reconstruction of the final refined map. **c-d**, Local resolution of the final refined map.

**Supplementary Figure 3.**
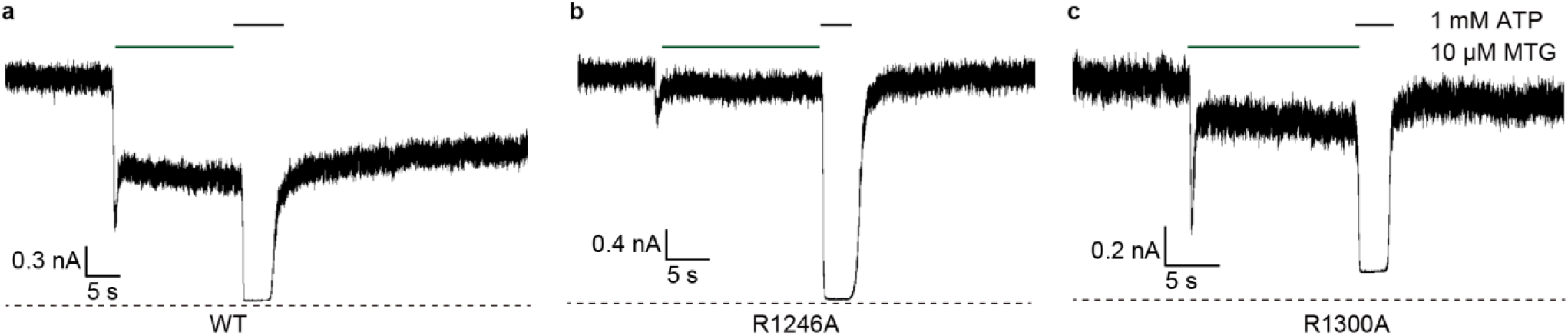
Representative recordings of MTG inhibition of K_ATP_ channels. **a-c**, The inhibition of 10 μM MTG inhibits on the inside-out currents of Wild-type, R1246A and R1300A K_ATP_ channels.

**Supplementary Table 1.**
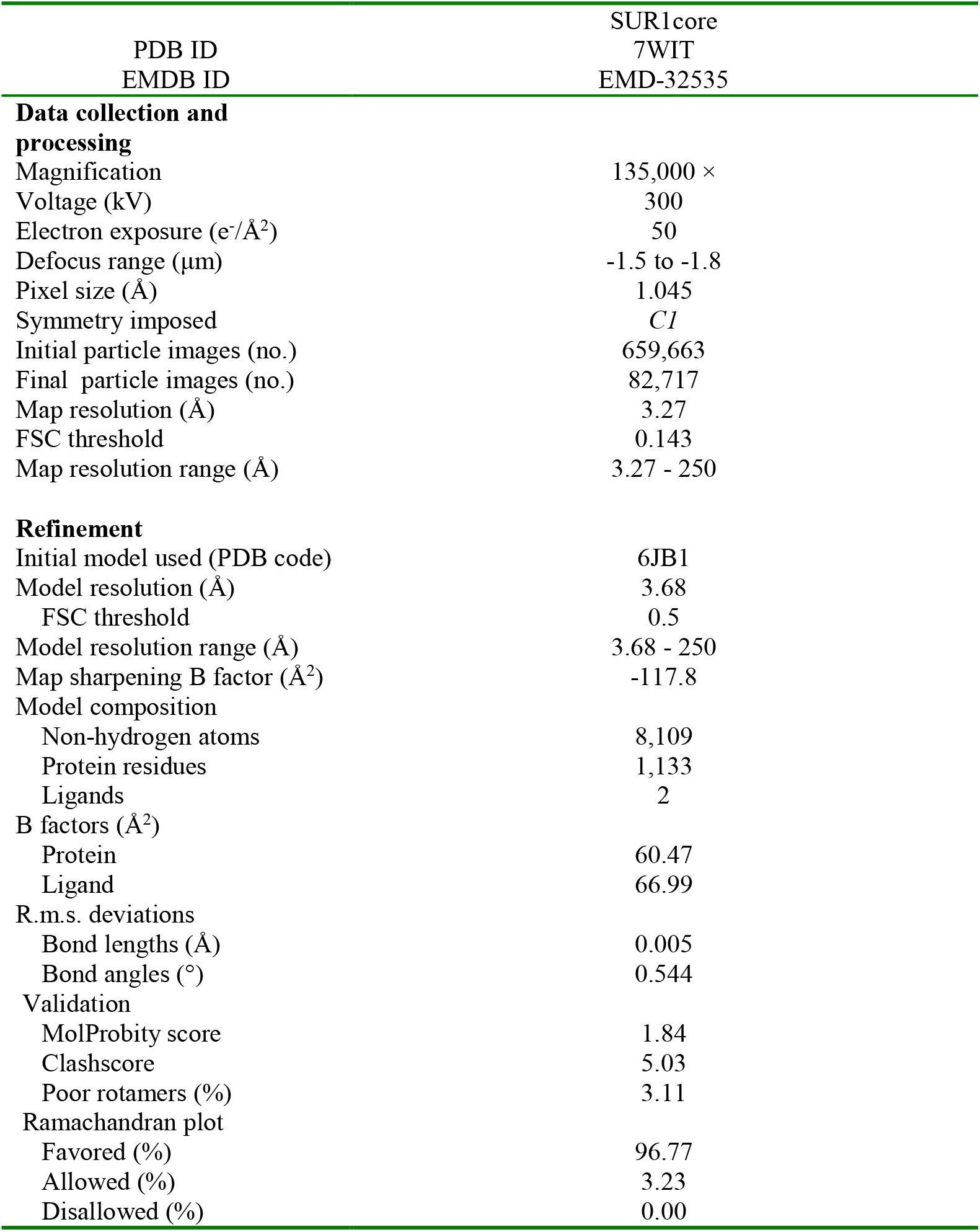
Cryo-EM data collection, refinement and validation statistics.

| PDB ID<br>EMDB ID | SUR1core<br>7WIT<br>EMD-32535 |
| --- | --- |
| **Data collection and processing** |  |
| Magnification | 135,000 × |
| Voltage (kV) | 300 |
| Electron exposure (e<sup>-</sup>/Å<sup>2</sup>) | 50 |
| Defocus range (μm) | -1.5 to -1.8 |
| Pixel size (Å) | 1.045 |
| Symmetry imposed | *C1* |
| Initial particle images (no.) | 659,663 |
| Final particle images (no.) | 82,717 |
| Map resolution (Å) | 3.27 |
| FSC threshold | 0.143 |
| Map resolution range (Å) | 3.27 - 250 |
| **Refinement** |  |
| Initial model used (PDB code) | 6JB1 |
| Model resolution (Å) | 3.68 |
| FSC threshold | 0.5 |
| Model resolution range (Å) | 3.68 - 250 |
| Map sharpening B factor (Å<sup>2</sup>) | -117.8 |
| Model composition |  |
| Non-hydrogen atoms | 8,109 |
| Protein residues | 1,133 |
| Ligands | 2 |
| B factors (Å<sup>2</sup>) |  |
| Protein | 60.47 |
| Ligand | 66.99 |
| R.m.s. deviations |  |
| Bond lengths (Å) | 0.005 |
| Bond angles (°) | 0.544 |
| Validation |  |
| MolProbity score | 1.84 |
| Clashscore | 5.03 |
| Poor rotamers (%) | 3.11 |
| Ramachandran plot |  |
| Favored (%) | 96.77 |
| Allowed (%) | 3.23 |
| Disallowed (%) | 0.00 |

